# Class imbalance should not throw you off balance: Choosing the right classifiers and performance metrics for brain decoding with imbalanced data

**DOI:** 10.1101/2022.07.18.500262

**Authors:** Philipp Thölke, Yorguin-Jose Mantilla-Ramos, Hamza Abdelhedi, Charlotte Maschke, Arthur Dehgan, Yann Harel, Anirudha Kemtur, Loubna Mekki Berrada, Myriam Sahraoui, Tammy Young, Antoine Bellemare Pépin, Clara El Khantour, Mathieu Landry, Annalisa Pascarella, Vanessa Hadid, Etienne Combrisson, Jordan O’Byrne, Karim Jerbi

## Abstract

Machine learning (ML) is increasingly used in cognitive, computational and clinical neuroscience. The reliable and efficient application of ML requires a sound understanding of its subtleties and limitations. Training ML models on datasets with imbalanced classes is a particularly common problem, and it can have severe consequences if not adequately addressed. With the neuroscience ML user in mind, this paper provides a didactic assessment of the class imbalance problem and illustrates its impact through systematic manipulation of data imbalance ratios in (i) simulated data and (ii) brain data recorded with electroencephalography (EEG) and magnetoencephalography (MEG). Our results illustrate how the widely-used Accuracy (Acc) metric, which measures the overall proportion of successful predictions, yields misleadingly high performances, as class imbalance increases. Because Acc weights the per-class ratios of correct predictions proportionally to class size, it largely disregards the performance on the minority class. A binary classification model that learns to systematically vote for the majority class will yield an artificially high decoding accuracy that directly reflects the imbalance between the two classes, rather than any genuine generalizable ability to discriminate between them. We show that other evaluation metrics such as the Area Under the Curve (AUC) of the Receiver Operating Characteristic (ROC), and the less common Balanced Accuracy (BAcc) metric – defined as the arithmetic mean between sensitivity and specificity, provide more reliable performance evaluations for imbalanced data. Our findings also highlight the robustness of Random Forest (RF), and the benefits of using stratified cross-validation and hyperprameter optimization to tackle data imbalance. Critically, for neuroscience ML applications that seek to minimize overall classification error, we recommend the routine use of BAcc, which in the specific case of balanced data is equivalent to using standard Acc, and readily extends to multi-class settings. Importantly, we present a list of recommendations for dealing with imbalanced data, as well as open-source code to allow the neuroscience community to replicate and extend our observations and explore alternative approaches to coping with imbalanced data.

## 1. Introduction

The rise of artificial intelligence (AI) in the last decade has led to important breakthroughs across many areas of science, including neuroscience and neuroimaging. New synergies between neuroscience and AI promise to drive both fields forward (1; 2; 3; 4; 5). In particular, machine learning is increasingly used both to model and to classify brain data (6), with applications ranging from cognitive and systems neuroscience (7) to clinical brain imaging (8; 9). As a result, machine learning is steadily turning into a fundamental tool for neuroscientists (10). As is the case with all methodological frameworks, machine learning comes with a set of subtleties and pitfalls. Being aware of these limitations and knowing how to handle them properly can be challenging, especially in research domains where machine learning is not yet adequately and systematically covered during training. The issue of data imbalance (11; 12) is a perfect example of an important problem that is generally well understood in the field of data science, but not always properly appreciated and tackled in neuroscience and neuroimaging. This technical note provides (1) a didactic description of the pitfalls associated with using skewed datasets in supervised machine learning, (2) a detailed assessment of the impact of varying the degree of class imbalance on classifier models and their performance using synthetic and real data, (3) concrete recommendations for mitigating the adverse effects of imbalanced data, and (4) open-source code to replicate the present work and extend it to other methods and metrics.

In binary classification problems, data imbalance occurs whenever the number of observations from one class (majority class) is higher than the number of observations from the other class (minority class)(11; 12). This problem is commonly encountered in cognitive neuroscience and in clinical applications, where observations for the target class (e.g. patients with neurological disorders) are often much harder to come by than for the control class (e.g. cognitively healthy individuals), leading to datasets with many more control observations than target observations (13; 11). Additional care has to be taken when evaluating the performance of diagnostic tests on rare conditions (14).

What makes imbalanced data problematic? When faced with highly skewed data, a classifier can achieve a high decoding accuracy merely by systematically and blindly voting for the majority class(11). For example, if an image classifier is asked to discriminate pictures of crows versus ravens, but only one out of twenty images in the training and test sets are ravens and the rest are crows, then the algorithm can (and likely will) achieve 95% accuracy simply by calling everything it sees a crow—though no discrimination or classification can rightly said to have been accomplished. In other words, given the opportunity, an algorithm will tend to bypass more complex feature analysis simply by “playing the odds”, which is indistinguishable from actual classification when only focusing on e.g. Accuracy as a performance metric.

The most common approach to avoid this problem is to enforce balanced data. One way to do this is by undersampling, i.e. by removing observations from the majority class until a balance is reached (15; 16), and repeating the process through bootstrapping. However, this comes at the cost of reducing the sample size, increasing the signal-to-noise ratio, which can be detrimental to the classification. Alternatively, one can oversample the minority class by duplicating or interpolating observations (17; 18; 19) (Fig. 1a), though this comes with a higher risk of overfitting and introducing noise (20).

**Figure 1:**
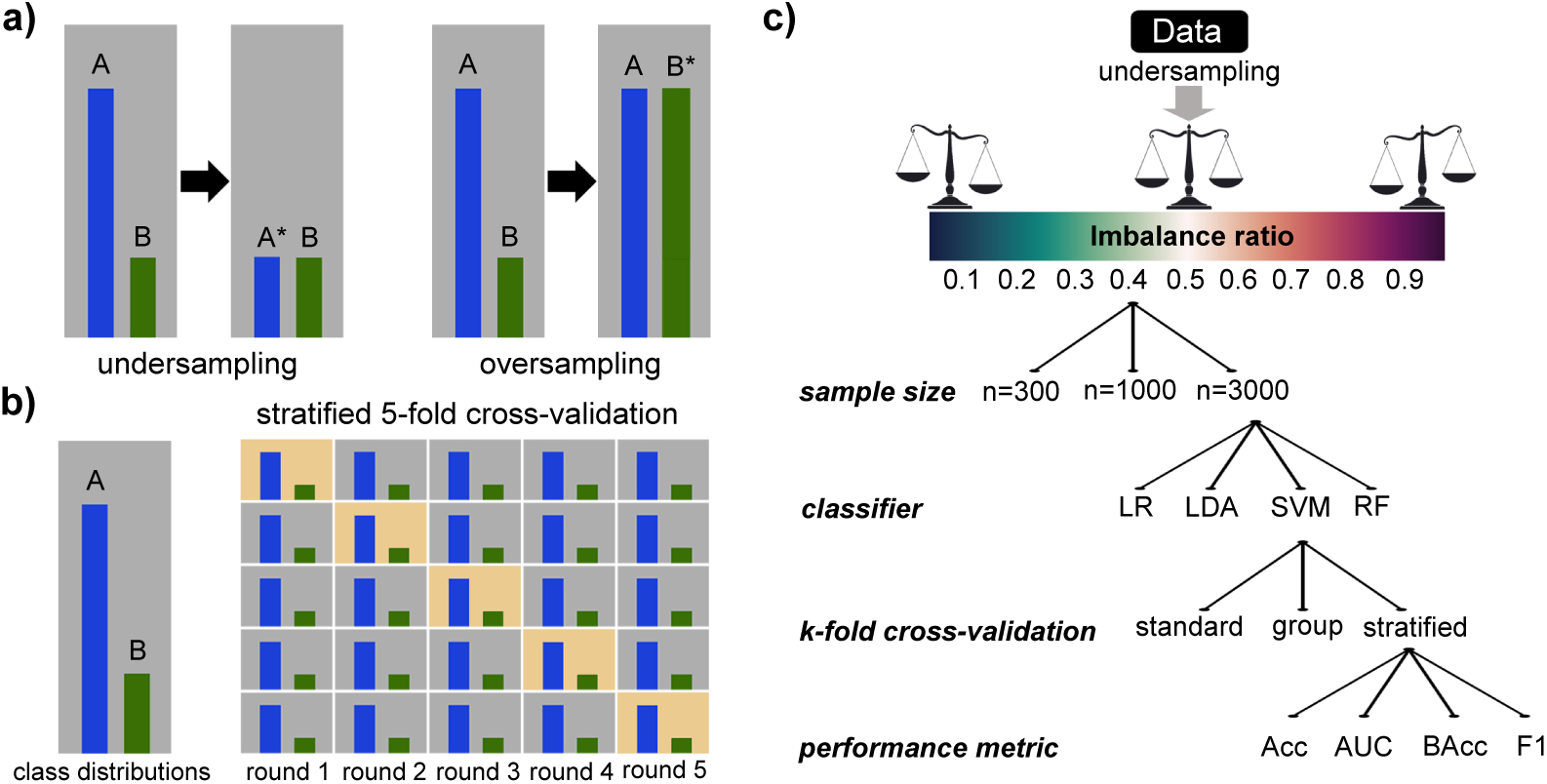
a) Methods to balance imbalanced data in order to avoid biases in machine learning. In undersampling (left), a subset of the overrepresented group (dataset A) is chosen. In oversampling (right), samples from the underrepresented group (dataset B) are duplicated or artificially augmented. **b)** Illustration of Stratified K-Fold cross validation (K=5). Instead of randomly choosing subsamples for every fold, this technique maintains the balance of the original data over all folds. This technique helps reduce biases and large variance in cross-validation. **c)** Illustration of the overall analysis framework of experiments performed in this paper. Various degrees of class imbalance were manually generated by undersampling the data. For a set of sample sizes, we performed binary classification using four widely used algorithms, three K-fold cross-validation methods, and four evaluation metrics (Acc, AUC, BAcc, and F1).

It may also be possible to dispense with undersampling or oversampling, and the problems they create, and to cope with the imbalanced data. In this case however, a number of additional considerations are necessary to avoid spurious results (21). These include the judicious choice of the type of classifier and the performance metric to be used. Additionally, when deploying a model validation scheme, special care must be taken to reflect the imbalance in the main data, such as by using Stratified K-Fold cross-validation (Fig. 1b). While these best practices are commonly applied in the machine learning community, they are not as widely adopted by the neuroscience and neuroimaging fields, likely due to the little information that exists, targeting neuroscientists, on how each of these different factors interact with data imbalance in a neuroscience context. Nevertheless, the importance of considering class imbalance has been highlighted in several brain decoding studies, and appropriate metrics have been made available in some toolboxes (22; 23; 24). Recommended metrics include the Area Under the Curve (AUC) of the Receiver Operating Characteristic (ROC), d-prime (a metric related to ROC-AUC, commonly used in psychology) (25; 26), balanced accuracy (27) and tuned loss functions (28). Yet, a dedicated account and systematic quantification of the effect of data imbalance for the neuroscience community is still missing; In particular, it is useful to have a didactic assessment of the impact of data imbalance (using both synthetic and real data) across varying degrees of imbalance, types of classifiers, hyperparameter choices, training, cross-validation and significance testing schemes. In this paper, we aim to provide a straightforward and practical demonstration of this multifaceted problem by using simulated data and real electrophysiological recordings. More specifically, we examine the behavior of four prominent metrics (Accuracy (Acc), Area Under the ROC Curve (AUC), Balanced Accuracy (BAcc) (29; 30), and F1; Table 1) across four widely-used classifiers (Logistic Regression (LR) (31), Linear Discriminant Analysis (LDA) (32), Support Vector Machine (SVM) (33), and Random Forest (RF) (34)), as we gradually increment data imbalance.

**Table 1:**
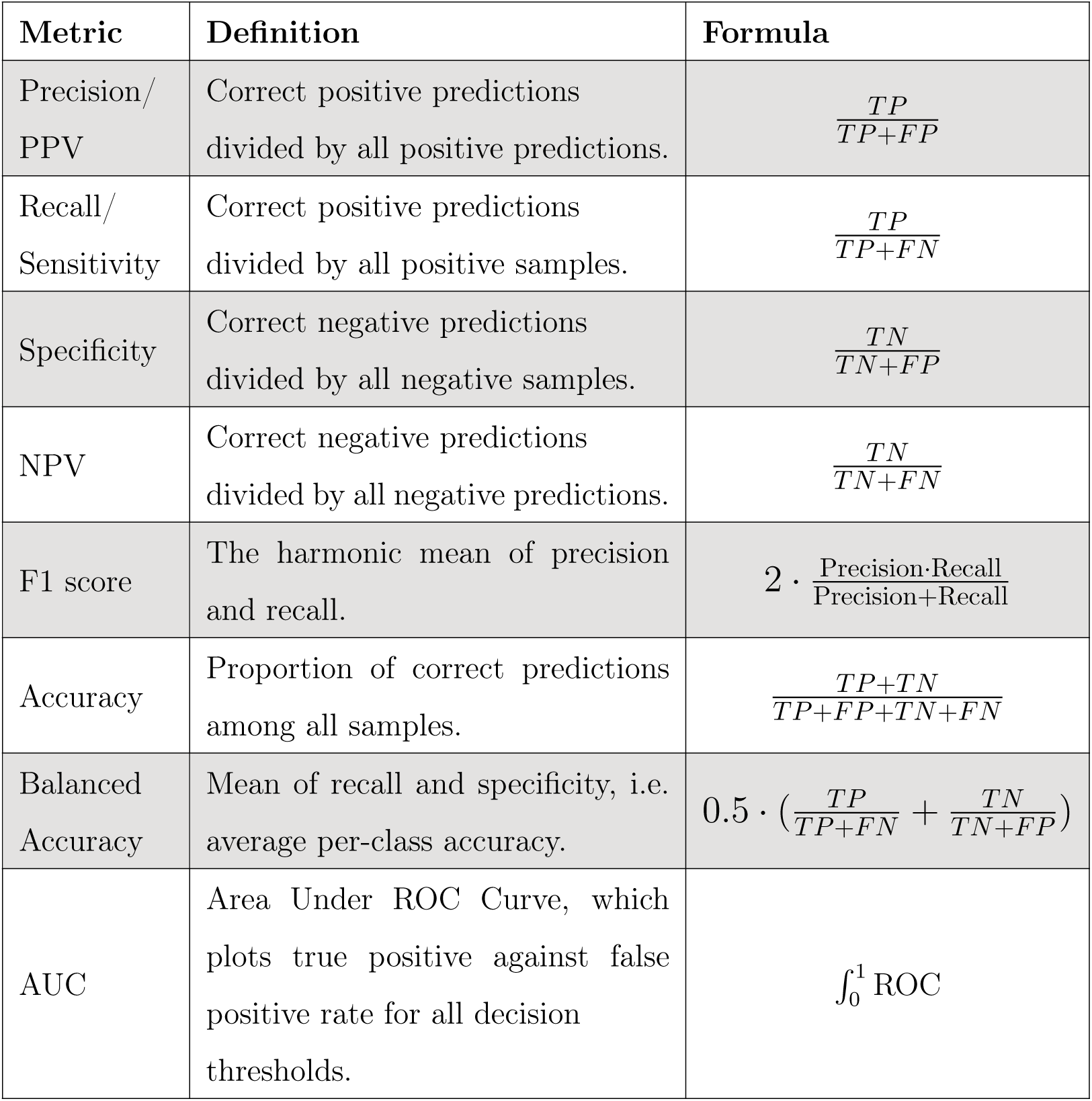
Overview of evaluation metrics. True positives (TP): instances that are positives and are classified as positives. False positive (FP): instances that are negatives and are classified as positives. False negative (FN): instances that are positives and are classified as negatives. True negatives (TN): instances that are negatives and are classified as negatives. PPV, Positive Predictive Value. NPV, Negative Predictive Value.

| Metric | Definition | Formula |
| --- | --- | --- |
| Precision/<br>PPV | Correct positive predictions<br>divided by all positive predictions. | $\frac{TP}{TP+FP}$ |
| Recall/<br>Sensitivity | Correct positive predictions<br>divided by all positive samples. | $\frac{TP}{TP+FN}$ |
| Specificity | Correct negative predictions<br>divided by all negative samples. | $\frac{TN}{TN+FP}$ |
| NPV | Correct negative predictions<br>divided by all negative predictions. | $\frac{TN}{TN+FN}$ |
| F1 score | The harmonic mean of precision<br>and recall. | $2 \cdot \frac{\text{Precision} \cdot \text{Recall}}{\text{Precision} + \text{Recall}}$ |
| Accuracy | Proportion of correct predictions<br>among all samples. | $\frac{TP+TN}{TP+FP+TN+FN}$ |
| Balanced<br>Accuracy | Mean of recall and specificity, i.e.<br>average per-class accuracy. | $0.5 \cdot \left( \frac{TP}{TP+FN} + \frac{TN}{TN+FP} \right)$ |
| AUC | Area Under ROC Curve, which<br>plots true positive against false<br>positive rate for all decision<br>thresholds. | $\int_0^1 \text{ROC}$ |

The topic of data imbalance, also often referred to as class or domain imbalance, has been addressed in previous work and online resources (11; 21), primarily within the computer science community. Here, we tailor our examples, explanations and recommendations, as well as our open-source code, to the neuroscience researcher or trainee with an interest in applying machine learning to electrophysiological data.

## 2. Methods and Materials

To explore the effect of data imbalance on different classification algorithms and performance metrics, we developed a custom open-source analysis pipeline, which systematically manipulates class imbalance (Fig. 1c). We herein first describe the analysis pipeline and secondary analysis, and then describe the five datasets used in this study (i.e. three types of simulated data, one EEG dataset, and one MEG dataset).

### 2.1. Primary Analysis Pipeline

The analysis pipeline was developed specifically for binary classification problems. Its primary purpose is to generate scores for different metrics across a range of imbalance ratios, using a list of classifiers and cross-validation schemes (Fig. 1c).

In order to estimate the chance level of correct classification, given the configuration of dataset and performance metric, the pipeline performs permutation tests (35) (repeatedly training and evaluating a classifier on the same dataset but with randomly permuted labels). Generating data-driven chance level is necessary because the theoretical binary classification chance level of 0.5 could be incorrect when performing binary classification on a dataset which has imbalanced classes or a small sample size. Furthermore, different performance metrics can lead to different estimates of the chance level.

In addition to that, we repeated experiments 10 times with different random seeds to estimate the degree of variance across cross-validation splits. We generally report the mean performance across the 10 repetitions and indicate the standard deviation as a shaded area around the mean.

To assess the impact of the metric with which classifiers are evaluated, we explored a range of classification metrics. These include Accuracy (Acc) (36), Balanced Accuracy (BAcc) (37), Area Under the ROC Curve (AUC) (38), and F1 (39). We made sure to include the most frequently used metrics, as well as variations specifically designed to tackle evaluation of prediction on imbalanced data. See Table 1 for an overview of classification metrics. It is interesting to note here that the Area under the Curve (AUC) of the Receiver Operating Characteristic (ROC) measures the integral of true positive rate against the false positive rate across all decision thresholds. This means that AUC is invariant to modifying the decision threshold, which is sometimes used to combat the influence of class imbalance.

In terms of classifiers, we included Logistic Regression (LR) (31), Linear Discriminant Analysis (LDA) (32), Support Vector Machine (SVM) (33), and Random Forest (RF) (34). We chose these models because they are among the most widely used in the neuroscience community, and because they represent a variety of approaches. The SVM was used with a radial basis function (RBF) kernel, making it a powerful non-linear classification algorithm. RF was of specific interest as it is a tree-based ensemble model expected to be better at handling class imbalance than the other methods.

We essentially used the default hyperparameters as defined in the scikit-learn library (40), however, we reduced the number of Random Forest estimators from 100 to 25 to better suit the low number of features in our experiments. Moreover, computing AUC requires predicting a continuous classification score for each sample. While LR, LDA and RF provide probability predictions out of the box, SVM supports two ways of estimating these prediction scores: probability calibration through an internal cross-validation on the training set, or using the signed distance from the fitted hyperplane. We found no substantial difference between both techniques and used the probability calibration method throughout the analysis. Note that AUC and Balanced Accuracy would be equivalent if we used binary predictions instead of class probabilities.

Cross-validation was performed using the Stratified K-fold or Stratified Group K-fold strategy (5 folds), depending on the presence or absence of group/subject information in the data. In a typical data-driven neuroscience decoding task, group labels help separate data from different subjects and add a measure of generalisation performance to new subjects to the evaluation process.

To simulate different amounts of imbalance in the class distribution we artificially limited the number of samples for both classes separately. We used a range of imbalance ratios from 0.1 (9:1 balance between the two classes) to 0.9 (1:9 balance) with 25 linearly spaced intermediate ratios, which provided a good trade-off between speed and performance. For a dataset with 100 data points (50 in either class), for example, we ran experiments with the following class distributions: 50:5, 50:7,…, 50:50,…, 7:50, 5:50. Imbalance was achieved by undersampling (dropping samples) either one of the two classes. It is important to note here that the sample size used to fit the classifiers decreased with increasing levels of imbalance. This limitation comes in part from the limited amount of data in our EEG and MEG analysis. We investigated the effect of sample size on our analysis in later stages of analysis, ensuring that it does not interfere with our results.

### 2.2. Secondary Analysis

Additionally, we explored the effect of data imbalance as a function of 1) the selection of hyperparameters, 2) the size of the dataset, 3) the type of cross-validation, 4) the effect of a balanced hold-out test set and 5) the impact of class imbalance on statistical significance testing. To simplify our approach, we only performed analysis 2-5 using SVM with an RBF kernel and evaluated it using Acc. We chose SVM specifically because we expected this algorithm to display important effects of class imbalance on performance.

#### 2.2.1. Effect of Hyperparameters

To assess the putative effect of hyperparameters, we explored those that are expected to have a significant impact on the robustness of classifiers with respect to imbalanced data (41). We used synthetic data (section 2.3) with 1000 samples (at perfect balance) and a distance of one between the two Gaussian distributions and evaluated the effect of the selected hyperparameters using Balanced Accuracy. This allows us to track improvements in robustness, which would manifest as a flatter curve of classification scores across imbalance ratios.

We limited this experiment to hyperparameters implemented in scikitlearn as this is one of the most commonly used libraries. Logistic Regression, SVM, and Random Forest all implement an automatic class-weighting algorithm to deal with imbalanced data, which can be enabled by setting the *class_weight* parameter to *balanced*. This approach weights the influence of each sample according to the inverse frequency of the corresponding class, thereby decreasing the impact of the majority class. This class-weighting technique, also known as cost-sensitive learning, penalizes the model less for errors made on examples from the majority class and more for errors made on the minority class. The Random Forest additionally has a *balanced_subsample* option, which applies the same weighting on the level of individual trees instead of globally for the full model. In addition to class weighting, we explored changing the minimum size of leaf nodes as a fraction of all samples (*min_weight_fraction_leaf*). As the fractional size of the leaf nodes depends on class distribution a large enough value ensures that leaf nodes will be more representative of differences between classes instead of simply voting for the majority class. We explored values of 0.1 and 0.4 for this parameter, serving as examples for weak and strong regularization.

While these hyperparameter optimization strategies are readily available for LR, SVM and RF in their respective scikit-learn implementations, tackling class imbalance by changing model parameters in the case of LDA is less straightforward. One simple strategy would be to modify the intercept of the decision hyperplane according to the rate of class imbalance. While this can be achieved through the *priors* hyperparameter in scikit-learn, exploration of this hyperparameter in more detail goes beyond the scope of this article. More generally, while hyperparameter optimization is not done systematically in brain decoding work, it is likely to become more common as ML continues to be increasingly used in neuroscience. Hence, understanding its impact on the issue of class imbalance could become increasingly relevant.

#### 2.2.2. Effect of Sample size

To test the impact of dataset size on robustness to imbalance we evaluated classifier performance across imbalance ratios using synthetic data with sample sizes N=300, N=1000 and N=3000 before unbalancing. The data of the two classes was sampled from two Gaussian distributions with a distance of one (section 2.3). This experiment further allows us to shed light on the issue of decreasing sample size with increasing levels of class imbalance, which results from the technique we used to generate imbalanced datasets.

#### 2.2.3. Effect of Cross-Validation

In order to assess the influence of the cross-validation scheme on different metrics when using imbalanced data, we tested K-Fold and Stratified K-Fold cross-validation on synthetic data. This difference will likely appear on smaller sample size since lower sample sizes increases the likeliness of having one class absent from a fold when using K-Fold without stratification.

To assess the impact of the choice of cross-validation approach, we trained an SVM classifier on the synthetic data with a distance of one between the means of both distributions (section 2.3) and chose to only use 50 samples per class before unbalancing. This analysis was repeated with 40 different seeds in order to assess the robustness of the effects we hypothesise.

#### 2.2.4. Effect of Balanced Hold-out Set

While so far all cross-validation splits came from the training data distribution, thereby replicating class imbalance, we also decided to explore the effects of training on an imbalanced dataset and performing validation on a balanced subset. Here, we trained an SVM on 1000 samples (at perfect balance) of synthetic data (section 2.3) with a distance of one between the means of both classes. The balanced hold-out set was created by taking a 10% split of the full dataset before artificially generating an imbalanced training set. This analysis aims at uncovering a potential performance bias when the train and test set have different class distribution, i.e. imbalanced and balanced respectively. Note that in this case, the Balanced Accuracy and Accuracy metrics are strictly equivalent as we are evaluating performance on a balanced hold-out set. We therefore only report accuracy.

#### 2.2.5. Significance testing on imbalanced data

We additionally evaluated the statistical significance of classification metrics across a range of imbalance ratios. Significance was computed from permutation tests with 100 permutations at p<0.01, using an SVM trained on 1000 samples (at perfect balance) of synthetic data with different amounts of overlap between classes, namely: identical distributions (impossible classification problem), a distance of 1 between the means of the class distributions (difficult classification problem) and a distance of 3 between the two classes (easy classification problem)(35).

### 2.3. Synthetic Data

To evaluate the impact of class imbalance in a controlled environment we generated synthetic data, consisting of 1000 random samples (at perfect balance) from two Gaussian distributions. We explored three different scenarios by modifying the amount of overlap between the two distributions, i.e. changing the distance between the means *µ*_1_ and *µ*_2_ of the two distributions, while keeping the standard deviation *σ*_1_ and *σ*_2_ constant at 1. In the first scenario, both classes came from the same distributions (*|µ*_0_ *− µ*_1_*|* = 0; Fig. 2a) and are therefore impossible to classify. In the second scenario, the two distributions were mostly overlapping (*|µ*_0_ *− µ*_1_*|* = 1; Fig. 2f), simulating a hard classification task. In the third scenario, the two distributions had a minimal overlap (*|µ*_0_ *− µ*_1_*|* = 3; Fig. 2k), which illustrates an easy classification task.

**Figure 2:**
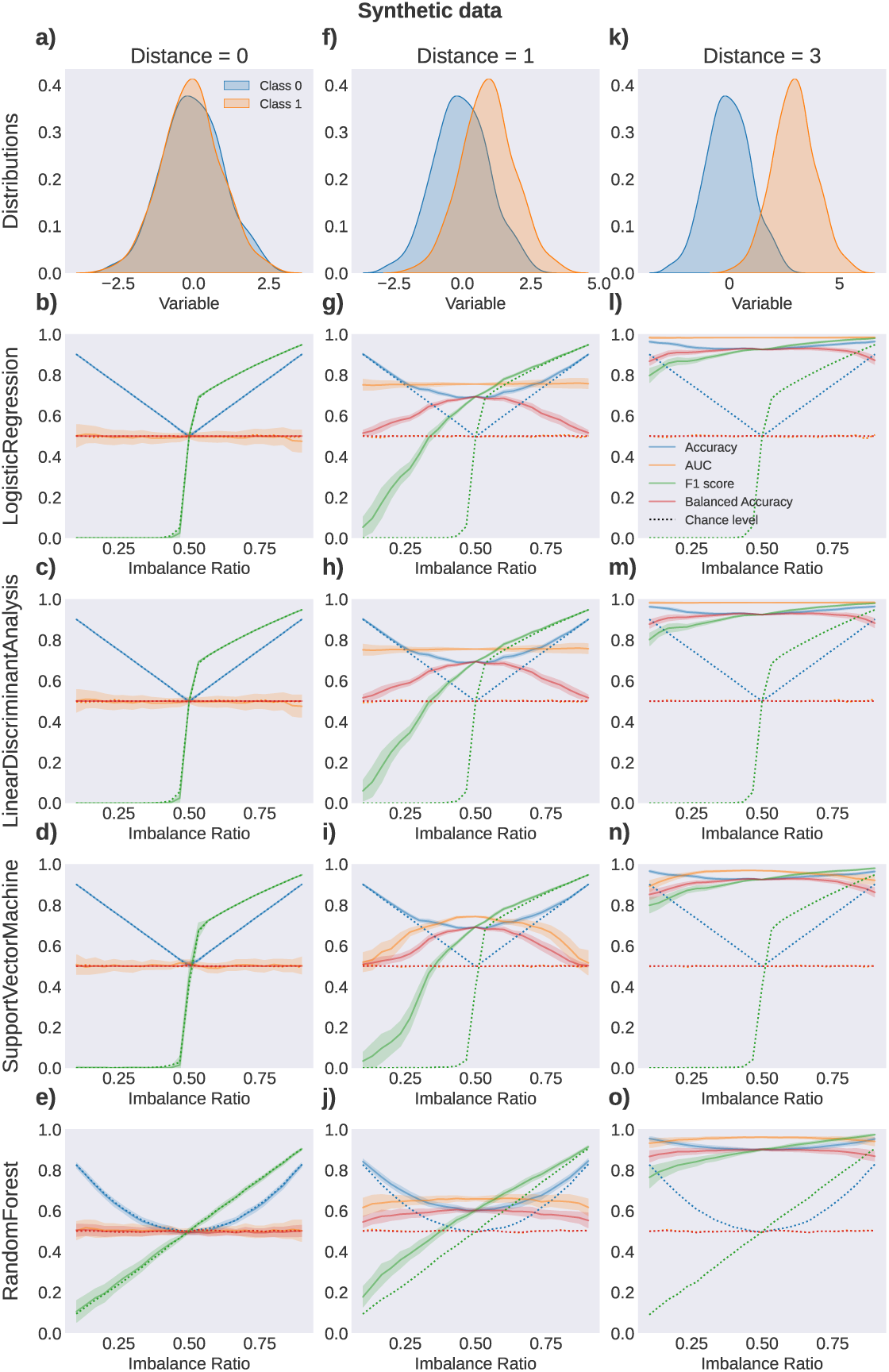
Effect of data imbalance on different performance metrics and algorithms using simulated data. This figure summarizes results from three different synthetic datasets (see class distributions in the first row): column 1 **(a-e)**: impossible classification task, all data comes from the same distribution, column 2 **(f-j)**: strongly overlapping class distributions and column 3 **(k-o)**: slightly overlapping class distributions. Rows 2 to 4 correspond to the different classification algorithms: Logistic Regression **(b,g,l)**, Linear Discriminant Analysis **(c,h,m)**, Support Vector Machine **(d,i,n)** and Random Forest **(e,j,o)**. We evaluated Accuracy (blue), AUC (orange), F1 score (_2_g_0_reen), and BAcc (red). Solid lines show the performance over different class imbalance ratios, averaged over 10 initializations. Colored areas represent the respective standard deviation. Dashed lines indicate the average performance over 100 random permutations (i.e. chance level) for every performance metric.

Note that this dataset serves a didactic purpose only, as we limit the simulated data to a single feature. While oftentimes one has to deal with multiple features, this example illustrates the problem at hand in a simplistic way and makes the concepts easy to grasp. In the following steps we extend our analysis to real-world examples using multiple features.

### 2.4. Brain Data

To extend the analysis from a controlled environment with synthetic data towards a realistic setting with electrophysiological datasets, we ran experiments on publicly available EEG and MEG datasets.

#### 2.4.1. EEG Motor Movement/Imagery Dataset

The publicly available EEG Motor Movement/Imagery Dataset (42; 43) consists of 64-channel EEG recordings of 109 subjects at 160Hz. While the dataset contains several tasks related to motor movement, only baseline resting-state runs were used, in this way creating a binary classification task between the eyes-open and eyes-closed conditions. Each recording has 1 minute of resting-state data which was segmented into 5-seconds epochs. As a result, 24 epochs per subject were extracted, half of them being eyes-closed and the others eyes-open. The effect of these conditions on neural oscillations is well studied, and consists of an increase in alpha power (8-12Hz) in posterior regions of the brain during eyes closed (44; 45), compared to the eyes open condition. We computed alpha power (8-12Hz) from the power spectral density (PSD) obtained using the multi-taper method. To restrict the features to the visual cortex, only parieto-occipital electrodes (17 out of the 64) were used. The total sample size of this dataset was 109 subjects *×* 2 conditions *×* 12 trials = 2616 samples at perfect balance.

A second analysis was carried out on this dataset to study the relationship between electrode locations and performance scores along 3 different imbalance ratios (0.1, 0.5 and 0.9). As topographic differences are the main focus of this experiment, the sensors were not constrained to be from parieto-occipital regions. Similarly, placing the emphasis on performance as a function of spatial location, single-channel, single-feature SVM classifiers were trained (in contrast to multi-feature classification in the previous analysis). The motivation behind this analysis is to tentatively illustrate that with increasing class imbalance, classifiers loose focus on the areas whose data best discriminate the classes, and merely predict the majority class.

#### 2.4.2. Cam-CAN Dataset

We used the passive auditory/visual perception task out of the open access MEG dataset collected at the Cambridge Centre for Ageing and Neuroscience (Cam-CAN) (46). The preprocessing steps for this dataset can be found in (47). The task consists of 2-min recordings during which subjects were presented with either visual checkerboards or auditory tones (in random order) 60 times each, with a second between each stimulus. We further processed the data by down-sampling to 500Hz and epoching into 800-millisecond trials with 150 milliseconds of signal before stimulus onset and 650 milliseconds after. The epochs were baseline-corrected before computing alpha power (12-30Hz) using the multi-taper method on the 650 milliseconds after onset. We excluded the magnetometers and averaged powers for the two gradiometers for each location. For this study, we randomly selected 20 subjects out of the 643 that are available in the repository, resulting in a sample size of 20 subjects *×* 60 stimuli *×* 2 stimulus types = 2400 samples at perfect balance. Classification was performed on the data of a single channel (Fig. 5b), which was selected by training separate classifiers for all channels and selecting the location with best performance.

### 2.5. Data and Code Availability

The scripts, notebooks and pipeline used in this study are open-source under the MIT licence. The code is available on GitHub for further explorations. Our experiment pipeline is not limited to the datasets explored in this study and can easily be used to explore other datasets. The open-source repository can be found at: https://github.com/thecocolab/data-imbalance.

The code was developed using Python and its rich ecosystem for scientific computing. To process brain data we used MNE-Python (48), and machine learning algorithms and metrics came from scikit-learn (40). Visualization was done with matplotlib (49) and seaborn (50).

The synthetic data used in this study can be generated using the open-source code we provide. The EEG Motor Movement/Imagery Dataset is publicly available and can be downloaded here: https://physionet.org/ content/eegmmidb/1.0.0/. The Cam-CAN dataset can be accessed upon request at https://camcan-archive.mrc-cbu.cam.ac.uk/dataaccess/ and the pipeline for preprocessing and loading the data is available at https://github.com/arthurdehgan/camcan.

## 3. Results

It is important to note here that it is not possible to compare absolute scores of metrics to one another. While for example AUC scores are generally higher than BAcc scores, this does not mean that the classifier performs better when using AUC. The performance metric does not influence the classifier’s behavior. In fact, we used the same fitted instance of a classifier to evaluate all performance metrics. The only exception here is that Acc and BAcc are mathematically equivalent in the case of perfectly balanced data. In the following we focus mostly on the shape of the graph across imbalance ratios in order to compare performance metrics.

### 3.1. Simulated data

To demonstrate the effect of data imbalance on different performance metrics, algorithms and cross-validation techniques, we simulated three binary-class datasets, as described in subsection 2.3 and Fig. 2a, f, k.

#### 3.1.1. Impossible classification

In the first scenario, the binary classification was performed on data from the same distribution (Fig. 2a-e), representing an impossible classification task. Due to the nature of the data, we expected performance values to stay at the chance level, which we estimated using random permutations (Section 2.1). This hypothesis was confirmed by our results, with all performance metrics staying close to their respective chance level (not the probabilistic chance level of 0.5; it varies as a function of data imbalance and metric etc.) and showing only minimal variation across repetitions. We herein first describe the effect of different performance metrics and then describe the difference between classification algorithms. AUC and BAcc showed identical behavior, staying consistently at a chance level of 0.5 across all levels of class imbalance. In contrast, Accuracy scores and the respective chance level increased towards both extremes of data imbalance and reached minimal values at the point of perfect data balance. More specifically, the Accuracy score consistently reflected the proportion of the majority class in the imbalanced data (i.e. reaching a score of 90 % for imbalance ratios of 9:1 and 1:9). The F1 score exhibited a steep increase from 0 towards 1 at the point where the data was balanced and continuously approached 1 with further increasing data imbalance.

Conspicuously, while Acc, AUC and BAcc show a symmetric pattern towards undersampling either class, the F1 score approaches 0 for increased undersampling of one class and 1 for the other. This behavior stems from F1 defining one class as positive and the other as negative, while the other evaluated metrics do not differentiate between the classes. F1 is a combined measure of the fraction of true positives among positive predictions (precision) and true positives among positive samples (recall). If the majority of samples are in the positive class, a classifier which always predicts the positive class will have precision and recall scores close to 1 and therefore high F1. On the other hand, an overrepresented negative class easily causes classifiers to always predict “negative”, resulting in an F1 score of 0. See (51) for a version of F1 adapted to imbalanced data.

It is noteworthy that the behavior of all above-described metrics was most similar between LR, LDA, and SVM, but varied in RF. Compared to the other algorithms, accuracy using RF exhibited a slower increase towards the extreme imbalance and stayed closer to 50% (i.e. the expected Accuracy score for classification between two datasets drawn from the same distribution) for a larger range of data imbalance ratios. In contrast to the steep increase of F1 described above, the F1 score in RF increased linearly over all levels of imbalance (Fig. 2b-e).

#### 3.1.2. Difficult classification

In the second scenario, binary classification was performed on data from two overlapping Gaussian distributions with a distance of one (Fig. 2f), representing a difficult classification task. Due to the nature of this dataset we expected performance scores to reach above chance level. Over all levels of data imbalance, AUC remained consistent and above chance level in LR and LDA. Only in SVM and RF did the AUC decrease to chance level on both ends of increased data imbalance, with the strongest decrease in SVM. Accuracy exhibited a similar behavior over all four classification algorithms. Similarly to the first scenario, the Accuracy scores reached maximal scores on both ends of data imbalance. Despite the decrease of Acc towards the point of maximally balanced data, the distance between Acc and its chance level was increasing. The maximum distance between Acc and corresponding chance level was reached at the point of perfect class balance, indicating that the classifier was best at learning the structure in the dataset at this point. In contrast to Acc, BAcc reached its maximum at maximal data balance and dropped to chance level, i.e. 50%, at both ends of class imbalance. Again, the RF showed more stable BAcc scores over the range of class imbalance. The F1 score successively increased from 0 to 1 and showed similar but dampened behavior compared to the results from the identical class distributions (Fig. 2g-j).

#### 3.1.3. Easy classification

In the third scenario, binary classification was performed on data from two Gaussian distributions with a distance of 3 (Fig. 2k), which is an example of an easy classification task. Due to the nature of the data, we expected performance values to reach high levels and to be less influenced by data imbalance. As expected, all performance metrics reached good classification scores which were less sensitive to data imbalance. While the general behavior was similar to the ones observed in the two previous experiments, the classifiers did a better job of learning structure from the data even in cases of imbalanced classes (Fig. 2l-o).

### 3.2. Secondary Analysis

#### 3.2.1. Hyperparameter Tuning

We explored the effect on robustness when tuning hyperparameters related to class imbalance of LR, SVM and RF. However, we only found improved robustness towards imbalance for LR by weighting samples inversely proportional to class frequency. While enabling this re-weighting scheme led to a stable BAcc score across imbalance ratios for LR (Fig. 3a), SVM, and RF remained vulnerable to class imbalance (Fig. 3b, c). We examined a second hyperparameter for the Random Forest, namely the minimum weight fraction (MWFL) to generate a leaf node (Fig. 3c). Increasing this hyperparameter beyond its default value of zero led to a general improvement in BAcc (over the default hyperparameter set) for balanced data, and a decrease towards regions of extreme class imbalance. Therefore, we are not able to report improvements in robustness from this hyperparameter for the Random Forest.

**Figure 3:**
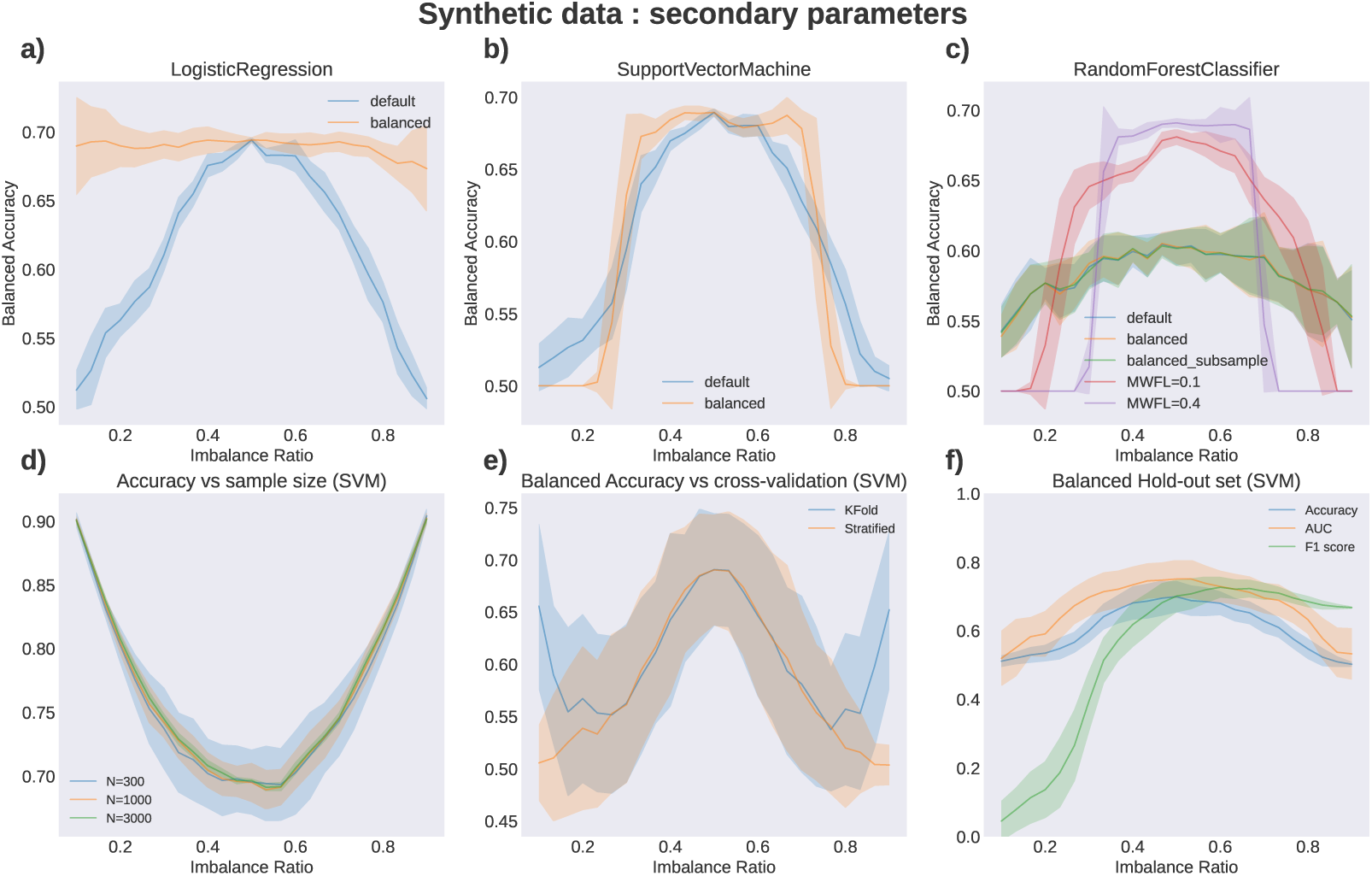
These examples follow the same principle as previous analysis (Fig 2), expanding on the effect of secondary parameters and hyperparameter tuning on the robustness to imbalance. **(a-c)** Exploration of the effect of hyperparameters on classifier robustness against imbalance. Each line represents a certain hyperparameter setting (Section 2.2.1) **d)** The effect of sample size on robustness to class imbalance using an SVM and Acc. **e)** The impact of cross-validation scheme on robustness to class imbalance (BAcc and SVM). **f)** Performance metrics on a balanced hold-out set using an SVM trained on different ratios of imbalance. BAcc is not shown here as it is equivalent to Acc for balanced data. Solid lines show the performance over different class imbalance ratios, averaged over 10 initializations. Colored areas represent the respective standard deviation.

#### 3.2.2. Sample Size

We did not find any notable effect of sample size (N=300, N=1000, N=3000) on overall SVM classification accuracy nor robustness to class imbalance. However, as one would expect, the variance across random initializations of the cross-validation splits decreased with the number of training samples (Fig. 3d). This result validates our procedure of undersampling to achieve class imbalance, which leads to a decrease in sample size towards higher degrees of imbalance. As here we show that a change in sample size only affects the variance of the results, the shape of the performance curves can be compared across imbalance ratios.

#### 3.2.3. Cross-Validation

Comparing different cross-validation algorithms revealed that K-Fold is more sensitive to imbalanced data than Stratified K-Fold (Fig. 3e). While both procedures led to similar Balanced Accuracy scores with balanced classes, K-Fold cross-validation without stratification showed an increase in Balanced Accuracy towards extremely imbalanced datasets. This likely stems from validation splits containing only a single class, which Balanced Accuracy does not account for. While a classifier voting for the majority class has a Balanced Accuracy of 0.5 on data containing two classes, its score will be 1 on validation splits that, by chance, only contain one class. The likelihood of this is higher for low sample sizes and is completely resolved by using Stratified K-Fold cross-validation.

#### 3.2.4. Balanced Hold-out Set

So far, all experiments were evaluated using cross-validation with imbalanced validation splits, replicating the class distribution of the training set. Figure 3f shows SVM classification scores on a balanced hold-out set after being trained on increasingly more imbalanced training data. While AUC and F1 scores are largely in line with previous results (Fig. 2i) using imbalanced cross-validation splits, Acc now reflects the previous behavior of BAcc, i.e. dropping towards the random baseline of 50% towards the extremes of class imbalance. Note that we do not report BAcc here as it is equivalent to Acc on balanced data.

#### 3.2.5. Significance testing on imbalanced data

Supplementary Figure 6 depicts significance scores across imbalance ratios and levels of difficulty of the classification problem. Generally we found that for the impossible classification task (i.e. identical class distributions; Fig. 6a), none of the scores were significant at p<0.01, as in this case, permuting labels does not remove any structure from the data. For the difficult classification task 6b), however, we found a range of statistically significant classification scores around perfect class balance. Scores were not significant towards the extremes of class imbalance. This behavior was shared among all the classification metrics we evaluated. In the third task—easy classification (Fig. 6c)—all classification scores were found to be significantly above chance level for all metrics, which highlights the classifier’s ability to learn structure in the data even for extreme class imbalance, which is even more pronounced for easier classification tasks. Note that the results in Figure 6 differ from the results presented in Figure 2d,i,n because — even though the experimental setup was the same — when testing for statistical significance we are limited to a single repetition of the analysis, while results in Figure 2 are averaged over 10 random seeds.

### 3.3. Results on EEG data

We performed two experiments using the EEG dataset with classification between eyes-open versus eyes-closed during resting (Section 2.4.1). We herein first describe the results of the multi-feature classification using parieto-occipital electrodes and then present the results of the channel-wise classification.

As previously shown in the simulated data, Acc increased with increasing data imbalance, reflecting the proportion of the majority class. In contrast to this, BAcc approached chance level with increased imbalance and reached its maximum with maximally balanced data. In line with the simulated data, the F1 score increased abruptly at optimal class balance. AUC was stable over a wide range of class imbalance. While SVM, LR, and LDA showed similar behavior (i.e. being equally sensitive to data imbalance), the performance metrics using RF exhibited more stability over different levels of imbalance (Fig. 4c-f).

**Figure 4:**
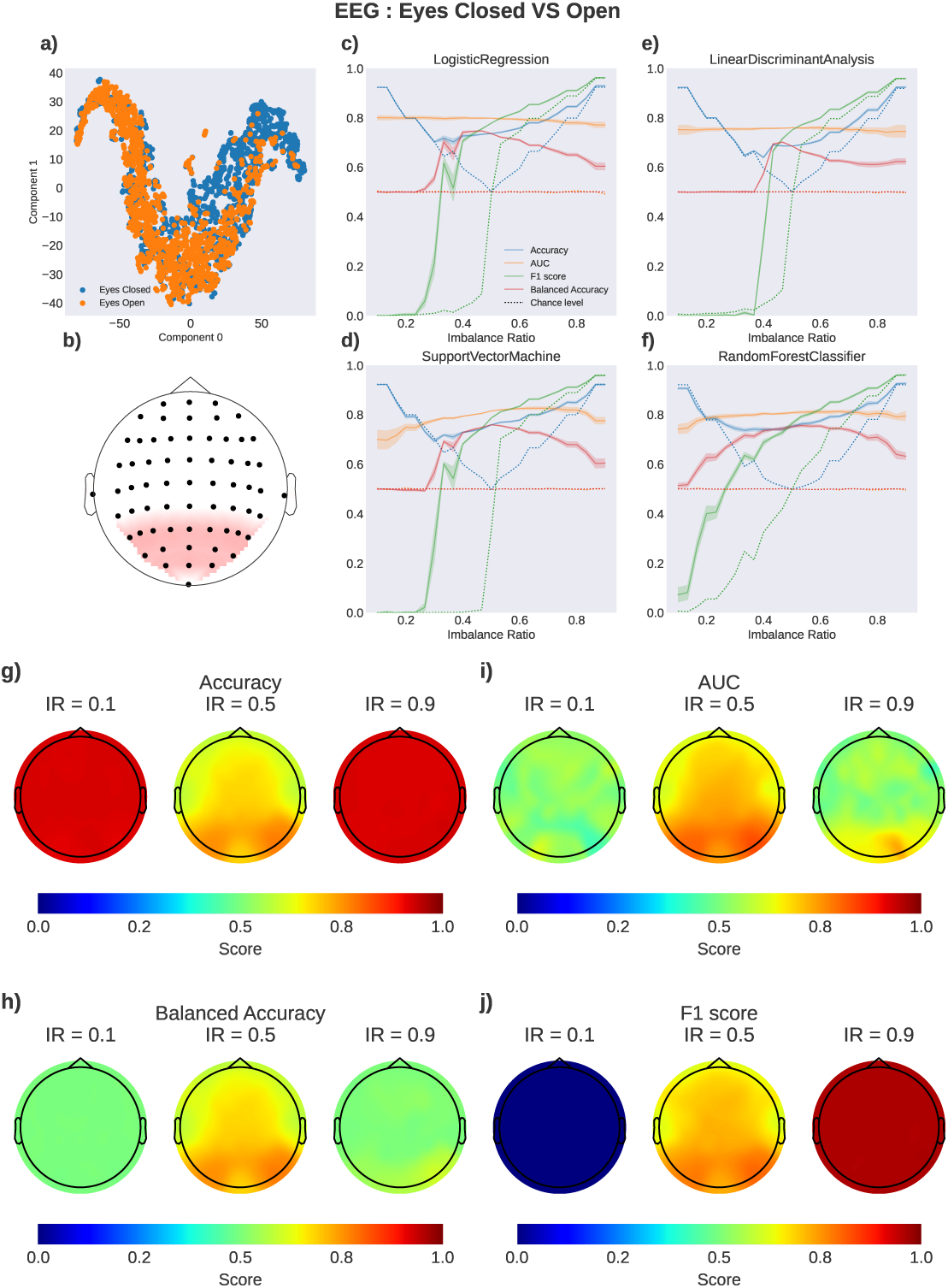
Effect of data imbalance on different performance metrics, algorithms, and cross-validation strategies for EEG data classification. **a)** 2D projection of the 17-dimensional input space using the t-SNE algorithm (52). Each dot represents one sample mapped onto two t-SNE components (x- and y-axis). This illustrates the amount of overlap (related to classification difficulty) between the eyes closed (blue) and eyes open (orange) classes. **b)** The parieto-occipital region of interest (ROI) was used as features. The effect of data imbalance using **c)** Logistic Regression; **d)** Support Vector Machine; **e)** Linear Discriminant Analysis, and **f)** Random Forest. We evaluated Accuracy (blue), AUC (orange), F1 score (green), and BAcc (red). Solid lines indicate performance across imbalance ratios averaged over 10 random initializations. Colored areas represent the respective standard deviation. Dotted lines indicate the average score acros_3_s_0_100 random permutations of class labels (i.e. data-driven chance level). Subfigures **g-j** highlight the effect between data imbalance and sensor location across performance metrics for single-channel, single-feature classification between eyes-open and eyes-closed EEG using an SVM. IR indicates the imbalance ratio used for each topographical map.

In contrast to multi-channel classification, single-channel decoding performance is commonly used to localize changes between two conditions and allows to attribute larger changes to channels with higher decoding performance (Fig. 4a, b). Here we show how class imbalance may lead to misinterpretation of such results and the loss of structure related to decoding performance. At optimal class balance, highest decoding performance was found in parieto-occipital regions of the brain, which is in line with the literature and allows for interpretation of the results (Fig. 4g-j, IR = 0.5). Towards the extremes of data imbalance, this structure is lost and we find uniform decoding performance across the brain. This effect is most prominent using Acc, BAcc, and F1, while AUC retains some variations across channels (Fig. 4g-j).

### 3.4. Results on MEG data

As described in Section 2.4.2, classification was performed between auditory and visual stimuli, using all sensor locations as features and averaged power across the two MEG gradiometers for each location. Figure 5 shows the results of the multi-feature classification using different models. All models seem to perform similarly and have similar reaction to imbalance with all of the studied performance metrics. The only notable difference in base-line chance level computations appears for the SVC classifier where F1 score raises sharply when reaching the balanced data ratio (Fig. 5). In line with the simulated and EEG datasets, BAcc approached chance level with increased imbalance and reached its maximum with perfectly balanced data. While AUC was stable across a wide range of imbalance ratios for LR, LDA, and RF, we observed comparable behavior to BAcc for SVM (i.e. approach-ing chance level towards more data imbalance). While SVM, LR, and LDA showed similar behavior (i.e. being equally sensitive to data imbalance), the performance metrics in RF exhibited slightly more stability over different levels of imbalance. This is in line with the aforementioned results using the simulated and EEG datasets.

**Figure 5:**
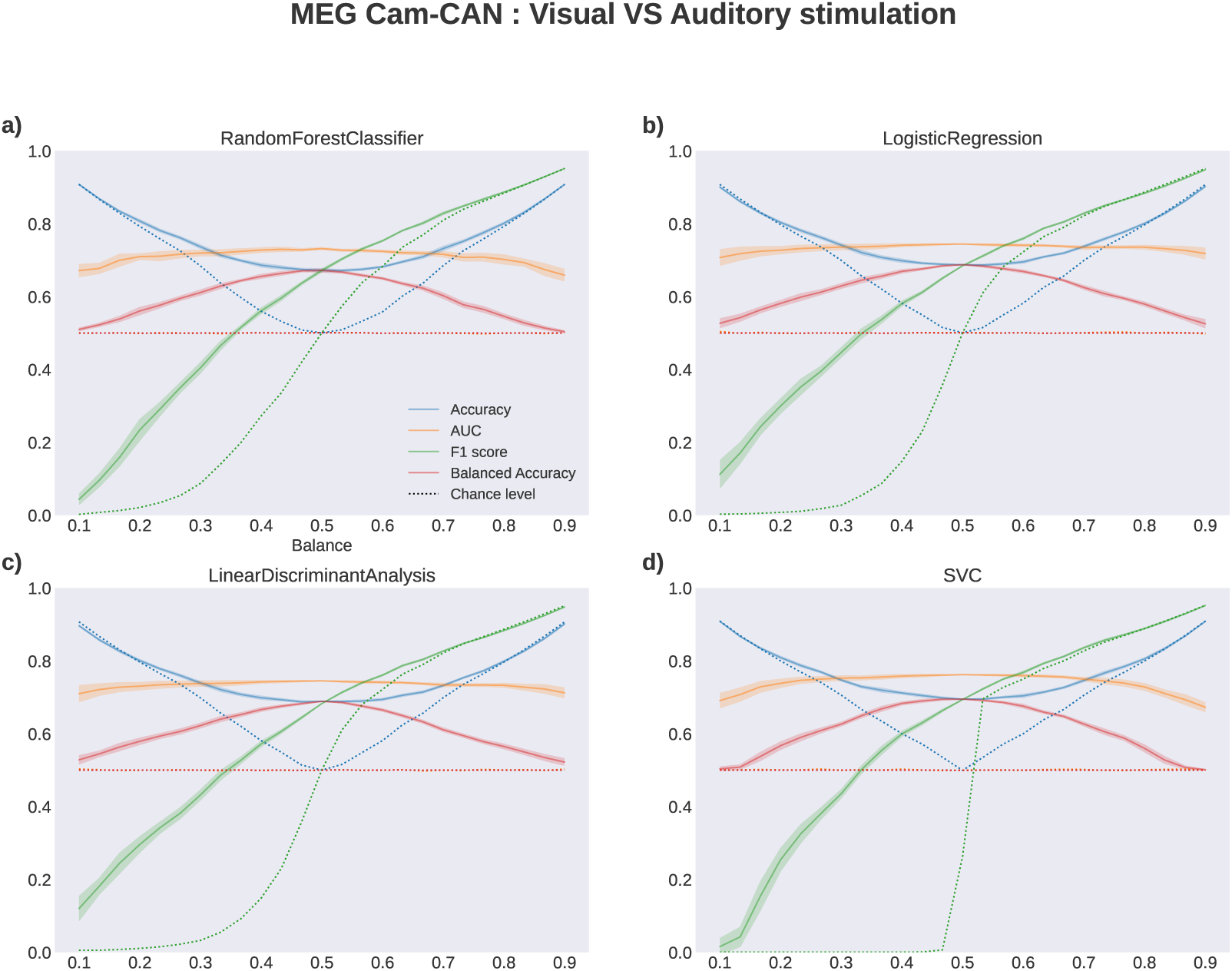
Effect of data imbalance on different performance metrics, algorithms, and cross-validation techniques on classification of MEG data. The effect of class imbalance using **a)** Logistic Regression; **b)** Random Forest; **c)** Linear Discriminant Analysis; **d)** Support Vector Machine. We evaluated Accuracy (blue), AUC (orange), F1 score (green), and BAcc (red). Solid lines indicate the performance over different class imbalances, averaged over 10 initializations. Colored areas represent the respective standard deviation. Dotted lines indicate the performance of 100 random permutations (i.e. chance level) for every performance metric.

## 4. Discussion

The present work shows that the implementation of classification on imbalanced data is feasible, though it demands certain important considerations. Our approach demonstrates how one needs to be mindful of class imbalance when choosing a classifier, an evaluation metric and a cross-validation scheme (13). Here, we sought to provide a didactic technical note on this question using a combination of simulated data and electrophysiological brain signals. Concretely, we quantified the behavior of commonly used classifiers, performance metrics, and cross-validation approaches across varying levels of data imbalance. An exhaustive exploration of all available techniques that have been proposed to tackle data imbalance is beyond the scope of this study. Instead, we chose to focus on machine learning tools and metrics that are often used within the neuroscience community. In line with this, the methods we address—and the open-source pipelines and notebooks we provide—all use the scikit-learn library.

Taken together, our observations support the idea that classification on moderately imbalanced data is feasible, as long as appropriate classifiers and performance metrics are employed. More specifically, by systematically manipulating the degree of data imbalance, we illustrated and quantified several key effects. First, we confirmed the tendency of classifiers to resort to blindly voting for the majority class as data imbalance was accentuated. When assessed with the widely used Accuracy measure, this behavior was associated with an artificial improvement in the model’s classification performance. AUC and BAcc were more robust to the increase in imbalance, and are therefore more appropriate under these circumstances. Moreover, our data confirms that Random Forest is more robust when handling imbalanced data, compared to other commonly used algorithms, such as LR, LDA, and SVM—especially when using class-weighting hyperparameter optimization. This result is expected, but our analyses quantify this for a wide range of imbalances and illustrates the effect with simulated data as well as EEG and MEG recordings. We also found that the balancing hyperparameter can be used to improve LR’s robustness to data imbalance.

Our study also highlights an important caveat concerning the use of permutation tests on imbalanced data. Permutation tests allow data-driven computation of the chance level, and from this chance level, they provide an estimate of statistical significance. However, because chance levels can be much greater than 50% in imbalanced data (for binary classification problems; Fig. 2, 6), a simple reporting in these cases of the Accuracy and of its statistical significance can artificially inflate the importance of the classification result. For example, for a 0.2 imbalance ratio, a statistically significant Accuracy of 82% can appear to be an outstanding result, when in reality, the chance level is 80%, so this is arguably comparable in importance to a statistically significant Accuracy of 52%. This scenario underlines the importance of reporting the chance level alongside performance metrics when carrying out permutation testing on imbalanced data (53).

The present study complements a wide array of insightful investigations that have explored the pitfalls and potential solutions for supervised learning with imbalanced data (11; 13; 54; 55; 56; 17; 57; 58; 39; 59; 20; 60; 14; 61). By contrast to some of the previous studies, the present work focuses on insights that are directly relevant to researchers in neuroimaging using standard tools, uch as those available through the scikit-learn library.

We would like to emphasize that while the examples provided here represent a specific instance of classification problems that can be encountered in the field of neuroscience, our analysis serves as an illustrative placeholder for many types of problems including but not limited to within-subject or between-subject classification, high dimensional problems as encountered with fMRI recordings, or analysis of behavioral data. Importantly, we encourage the reader to follow the same procedure to explore the imbalance question in their own data. As a matter of fact, the code and Jupyter notebooks (https://github.com/thecocolab/data-imbalance) were designed so the figures could be easily replicable, and allow users to extend the investigations to a wider range of metrics and methods, in a collaborative and open science perspective.

### Recommendations

In discussing recommendations and best practices for handling data imbalance, it is important to note that the utility or suitability of a metric should always be determined in relation to the specific problem being tackled. The various types of machine learning problems differ among other things in the type of error one seeks to minimize—this will in turn determine the appropriate classification paradigm in which to operate. The suitable paradigm depends on whether one’s aim is to minimize overall classification error (62), type I error (false positive) (63) or type II error (false negative) (64). Of these three, the design of the present study and the findings we report relate primarily to paradigms seeking to minimize overall classification error. Our analysis confirms that BAcc (and the related AUC metric) are more robust to data imbalance than the common Accuracy metric. Note that the Precision and Recall metrics are recommended when dealing with type I and type II errors respectively. Sampling techniques (i.e. over- and undersampling) are known to be helpful in most paradigms and evidence suggests that they work well in combination with certain classifiers(65).

Based on the observations reported in this study, evaluating the performance metrics on a balanced holdout set (Fig. 3f) allows for an unbiased evaluation of classification performance even when using metrics vulnerable to class imbalance, such as Acc. We therefore recommend using several evaluation metrics, e.g. BAcc, with Stratified K-Fold cross-validation and standard Accuracy on a balanced holdout set. As BAcc is equivalent to Acc for balanced data, it retains Acc’s greatest advantage, namely its intuitiveness and ease of interpretation. We additionally argue that BAcc results in more intuitive performance evaluation for imbalanced data, as it combines performances of individual classes with equal weight. Accuracy on the other hand combines class performances with a strong bias towards the majority class.

Deep learning, an advanced type of machine learning used for a large variety of classification tasks, is not immune to data imbalance (66). Deep learning models learn by backpropagating gradients through the model. In class-imbalanced scenarios, the majority class dominates the net gradient that is responsible for updating the model’s weights, which reduces the error of the majority group quickly during early iterations. However, oftentimes it simultaneously increases the error of the minority group. As a result, the neural network struggles to learn the decision boundary for the problem (67). Common approaches used to overcome data imbalance when training deep neural networks include under-/oversampling, data augmentation, the use of a class-weighted loss function (i.e. higher penalty for errors made on the minority class) and output thresholding (68).

Overall, given our results, we make the following seven recommendations for machine learning in neuroscience:

1. **Know your problem.** The right performance metric to use is determined first and foremost by the specific research question at hand. Throughout this study, we have focused largely on the type of question that requires a minimization of the overall classification error. However, for some problems, it may be more important to prioritize the minimization of either type I or type II error. In these cases, other performance metrics may be more relevant than BAcc or AUC, namely Precision and Recall. For example, a classifier used to discriminate biological sex based on brain activity would likely seek to minimize overall classification error, and BAcc or AUC would then be recommended to correct for any class imbalance in the data. In contrast, a classifier used to detect malignant tumours in brain imaging would instead benefit most from Precision and Recall metrics, since false negatives (type II error) would need to be minimized above all else. Therefore, consideration of the nature of the problem at hand—specifically, of the type of error to be minimized—is essential to selecting the most appropriate performance metric. With this in mind, the remaining recommendations offered here apply specifically to problems requiring the minimization of classification error.
2. **Use Balanced Accurary (BAcc).** BAcc has been largely underexploited in neuroscience research. Given (i) its superior robustness to imbalance, (ii) the fact that it simply reduces to Accuracy for balanced datasets and (iii) its applicability to both binary and multi-class (69) datasets, we recommend the routine use of BAcc, rather than the commonly used Acc, as a default for neuroscience machine learning applications where overall classification error should be minimized. To maximize the interpretability of classification results however, it is worth looking at multiple performance metrics (e.g. BAcc and AUC). If the classifier is purely used to decide if a feature captures a difference between two conditions (as commonly done in the field of neuroscience), AUC combined with significance testing serves as a powerful tool.
3. **Use ensemble methods.** If data imbalance cannot be avoided, we recommend the use of classifier families that provide additional robustness. In line with previous recommendations, ensemble methods such as Random Forests are less sensitive to data imbalance and provide a set of hyperparameters that can be optimized for the classification of imbalanced data. Further, ensemble methods are generally known to improve robustness (70; 71). However, the complexity of some of these algorithms might not fit well onto very simple classification tasks, which may affect the robustness.
4. **Use a balanced hold-out test set.** When working with imbalanced data, the true overall classification error of the trained classifier can be assessed simply by testing it on a balanced hold-out test set. However, we further suggest also evaluating classifier performance on a test set that reflects the class distribution of the specific problem in the wild (or of the training set, if the class distribution in the real-life setting cannot be estimated). The difference in score between these two test sets can additionally help interpret the performance of the classifier. Beyond that, analysing confusion matrices and their derived metrics helps with understanding the behavior of classifiers in more detail and shed light on class imbalance related biases. For an illustration of this see (72).
5. **Use Stratified K-fold for cross-validation.** K-fold cross validation is highly sensitive to data imbalance such that in extreme cases, it can even become impossible to perform classification (i.e. if one fold contains only a single class). We therefore strongly recommend the use of Stratified K-fold, which maintains the imbalance ratio within each of the selected folds.
6. **Report statistical significance AND chance level.** Without the corresponding chance level, performance metrics can easily be misinterpreted. Especially for performance metrics like Acc, chance level fluctuates widely with data imbalance. Thus, performance scores should always be reported accompanied by—and should interpreted in the light of—the associated chance level. The chance level can be estimated in a data-driven approach using permutation tests. This recommendation also applies if statistical tests were performed and performance scores reached significance over random permutations.
7. **Use hyperparameters.** For many of its classifiers, scikit-learn provides hyperparameter options specifically designed for dealing with imbalanced data; these should be routinely exploited. Our study highlights the potential utility of this step and encourages the reader to consider hyperparameters for an optimal performance. An extensive tutorial on hyperparameter selection is beyond the scope of this work. There is substantial useful literature on hyperparameter selection for further reading (73; 74; 75; 76; 77; 78; 28).

### Limitations and perspectives

This study’s results need to be interpreted in the light of several limitations. First, many approaches for handling imbalanced data have been proposed (21). In this study, we focused on families of models and performance metrics that are easily accessible and widely used in the neuroscience community, in particular through the scikit-learn library. Specifically, we focused on four popular classifiers and four standard evaluation metrics, as well as two cross-validation schemes. Reviewing or comparing all existing tools is beyond the intended goal of this paper, but the Python code we provide is open-source, which allows users to extend these investigations. Second, the data we used consisted of simulated Gaussian data distributions, as well as open-access electrophysiological brain signals (MEG and EEG). We did not examine the impact of noise, though it could have been interesting to consider it, as it has been shown to interact with the performance of the classifier (79). Third, we only briefly mention the option of balancing the data through over and under-sampling methods, and focused our investigation on evaluating the impact of data imbalance. Fourth, the aim of this study was to explore the individual classifiers’ sensitivities and relative changes of performance metrics over systematically imbalanced data. Therefore, the presented results do not allow a comparison of individual classifiers’ absolute performance.

Fifth, our main focus was to explore the effect of data imbalance on classification tasks. To this end, we explored the simplest form, namely a binary classification, where the manipulation of the imbalance ratio is straightforward. While creating imbalanced data for the multiclass case is more complicated, most of our findings extend to this type of classification—with some caveats. For instance, AUC is typically available only for binary classification, although extensions for multiclass classification exist (80). In addition, some algorithms such as SVM and LR reframe multi-class classification as a separate one-vs-the-rest binary classification problem for each class; this effectively results in class imbalance. While other options exist, such as multinomial logistic loss (i.e. cross-entropy loss) for LR or one-vs-one classification for SVMs, we advise caution when using these algorithms in a multi-class setting. Notwithstanding, the main take-home message of this study—that BAcc is generally more robust than Acc—still applies for multiclass classification.

### Conclusion

In this study, we have illustrated the effect of imbalanced data on some of the most prominent classification algorithms and performance metrics used in neuroscience. Among other things, one key take-home message is our suggestion to systematically use Balanced Accuracy over the widely used Accuracy metric, whenever the aim is to minimize overall classification error. In addition to its robustness to class imbalance, Balanced Accuracy collapses to standard Accuracy for balanced datasets and is readily extendable to multiclass problems. More generally, we hope that the recommendations and red flags reported here—using simulations and real brain data—will strengthen good practices for the application of supervised ML in neuroscience, and increase awareness, especially among new-comers to the field. Lastly, by providing open-source code and well-documented pipelines, we hope that others will further explore this question with a wider variety of parameters, classifiers, and different types of classification problems.

## Acknowledgements

KJ was supported by funding from the Canada Research Chairs program and a Discovery Grant from the Natural Sciences and Engineering Research Council of Canada (NSERC), and IVADOApogée fundamental research project grant. YH was supported by the Courtois-Neuromod Project. CM was supported by the FRQNT Strategic Clusters Program (2020-RS4-265502 - Centre UNIQUE - Union Neurosciences and Artificial Intelligence – Quebec and the Canada First Research Excellence Fund and Fonds de recherche du Quebec, awarded to the Healthy Brains, Healthy Lives initiative at McGill University. HA was supported by Mitacs Globalink Research Award. JOB was supported by the Canadian Institutes of Health Research (CIHR). AD was supported by Mila. AK was supported by Mitacs Graduate Fellowship and Courtois foundation. LMB was supported by IVADO. PT acknowledges support through a scholarship from the Cognitive and Computational Neuroscience Laboratory (CoCo Lab) and Mila (Quebec Machine Learning Institute). TY was supported by Mitacs. YM was supported by Mitacs through the Mitacs Globalink Research Internship program. The authors wish to thank Kris, Mark, T. Hortons and Mr. and Mrs. Puffs for their support during the Hackathon/Paper Sprint events.

## Supplementary Material

**Figure 6:**
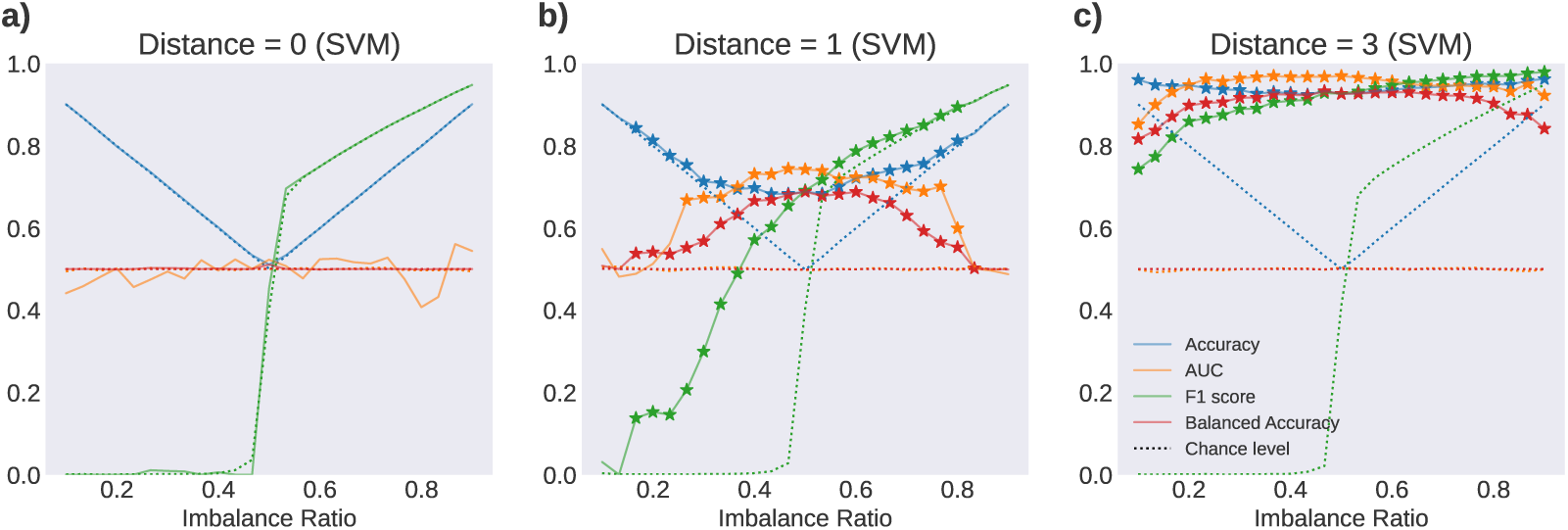
Statistical significance of different amounts of data imbalance and difficulty of the classification task, using an SVM on 1000 samples of synthetic data. Stars represent statistical significance at p<0.01, computed with permutation tests (N=100). **a)** Identical distribution for both classes (impossible classification problem). **b)** Hard classification problem with a distance of 1 between means of class distributions. **c)** The distance between the class distributions is 3, corresponding to an easy classification problem.

## Notes

https://github.com/thecocolab/data-imbalance

### Competing Interest Statement

The authors have declared no competing interest.

### Summary of Updates

clarifications based on reviewer's comments; adding citations to several claims; updating figures according to reviewer's comments

